# *Candida albicans* promotes neutrophil extracellular trap formation and leukotoxic hypercitrullination via the peptide toxin candidalysin

**DOI:** 10.1101/2022.10.13.512027

**Authors:** Lucas Unger, Emelie Backman, Borko Amulic, Fernando M. Ponce-Garcia, Sujan Yellagunda, Renate Krüger, Horst von Bernuth, Johan Bylund, Bernhard Hube, Julian R. Naglik, Constantin F. Urban

**Affiliations:** Department of Clinical Microbiology, Umeå University, Umeå, Sweden; Umeå Centre for Microbial Research (UCMR), Umeå University, Umeå, Sweden; School of Cellular and Molecular Medicine, University of Bristol, Bristol, UK; Department of Pediatric Respiratory Medicine, Immunology and Critical Care Medicine, Charité – Universitätsmedizin Berlin, Berlin, Germany; Labor Berlin Labor Berlin – Charité Vivantes GmbH, Department of Immunology, Berlin, Germany; Berlin Institute of Health at Charité – Universitätsmedizin Berlin, Germany; Charité - Universitätsmedizin Berlin, corporate member of Freie Universität Berlin, Humboldt-Universität zu Berlin, and Berlin Institute of Health (BIH), Berlin-Brandenburg Center for Regenerative Therapies (BCRT), Berlin, Germany; Department of Oral Microbiology & Immunology, Institute of Odontology, Sahlgrenska academy at University of Gothenburg, Gothenburg, Sweden; Department of Microbial Pathogenicity Mechanisms, Leibniz Institute for Natural Product Research and Infection Biology - Hans-Knoell-Institute, Jena, Germany; Friedrich Schiller University, Jena, Germany; Centre for Host-Microbiome Interactions, Faculty of Dentistry, Oral & Craniofacial Sciences, King’s College London, London, United Kingdom

**Keywords:** *Candida albicans*, candidalysin, PAD4 citrullination, ROS, NETs, chronic granulomatous disease, fungal immunology

## Abstract

The cytolytic peptide toxin candidalysin is secreted by the invasive, hyphal form of the human fungal pathogen, *Candida albicans*. Candidalysin is essential for inducing host cell damage during mucosal and systemic *C. albicans* infections, resulting in neutrophil recruitment. Neutrophil influx to *C. albicans*-infected tissue is critical for limiting fungal growth and preventing the fungal dissemination. Here, we demonstrate that candidalysin secreted by hyphae promotes the stimulation of neutrophil extracellular traps (NETs), while synthetic candidalysin triggers a distinct mechanism for NET-like structures (NLS), which are more compact and less fibrous than canonical NETs. Candidalysin activates NADPH oxidase and calcium influx, with both processes contributing to morphological changes in neutrophils resulting in NLS formation. NLS are induced by leukotoxic hypercitrullination, which is governed by protein arginine deaminase 4 activation via calcium influx and initiation of intracellular signalling events. However, activation of signalling by candidalysin does not suffice to trigger downstream events essential for NET formation, as demonstrated by lack of lamin A/C phosphorylation, an event required for activation of cyclin-dependent kinases that are crucial for NET release. Interestingly, exposure to candidalysin does not immediately restrict the capability of neutrophils to produce reactive oxygen species (ROS), nor to phagocytose particles. Instead, candidalysin triggers ROS production, calcium influx and subsequent activation of downstream signalling that drive morphological alteration and the formation of NLS in a dose- and time-dependent manner. Notably, candidalysin-triggered NLS demonstrate anti-*Candida* activity, which is resistant to nuclease treatment and dependent on the deprivation of Zn^2+^. This study reveals that *C. albicans* hyphae releasing candidalysin concurrently trigger canonical NETs and NLS, which together form a fibrous sticky network that entangles *C. albicans* hyphae and inhibits their growth. Importantly, this explains discrepancies of previous studies demonstrating that neutrophil-derived extracellular chromatin structures triggered by *C. albicans* can be both dependent and independent of ROS. Our data also demonstrate that while candidalysin hampers neutrophil function, the toxin also increases the capability of neutrophils to entangle hyphae and to restrict their growth, reflecting the importance of human neutrophils in controlling the dissemination of *C. albicans*.

## Introduction

Neutrophils are important innate immune cells that play a pivotal role in preventing fungal infections ^1^. In addition to engulfing and eradicating microbes by phagocytosis, extracellular mechanisms involving the release of neutrophil extracellular traps (NETs) and granular vesicles have been described ^2–4^. As pathogenic fungi can grow as a network of filamentous hyphae, phagocytic killing by neutrophils is often insufficient, thus extracellular mechanisms, such as NET formation, are required for efficient eradication. NETs have been reported to restrict fungal growth and to corroborate inflammatory responses during mycoses ^1,5,6^. Pathogenic fungi trigger NETs in an NADPH oxidase-dependent manner involving activation of cyclin-dependent kinases 4 and 6 (CDK4/6) ^7^. Several studies indicate that if NET release is not properly balanced, NETs may also have harmful effects on the host, mainly due to their pro-inflammatory function ^8^. Notably, microbial toxins can trigger leukotoxic hypercitrullination of histones in neutrophils resulting in similar extracellular structures, termed NET-like structures (NLS) ^9,10^. NLS are less fibrous and more compact than canonical NETs and are triggered in an NADPH oxidase-independent fashion. Similiar to NETs, NLS can induce pro-inflammatory effects with potentially hazardous consequences for the host ^10–12^.

The human fungal pathogen, *Candida albicans*, is a dimorphic yeast with the ability to form invasive, filamentous hyphae ^13^. The yeast-hyphal transition, in combination with the expression of hypha-associated factors, is critical for *C. albicans* virulence ^14,15^. Invasive *C. albicans* hyphae are controlled by human neutrophils, thereby preventing dissemination and exacerbation of disease in otherwise healthy patients ^16^. A critical factor for the invasive and inflammatory potential of *C. albicans* hyphae is the recently discovered peptide toxin candidalysin ^17,18^. Candidalysin is released from the polyprotein Ece1p via a sequential proteolytic cleavage by the proteases Kex2p and Kex1p ^19^. The corresponding *ECE1* gene is exclusively expressed by the hyphal morphology of *C. albicans* ^20^ and belongs to the hyphal core response genes consisting of eight hyphal-associated genes expressed under a variety of hyphal inducing conditions ^21^. *C. albicans* hyphae deficient in candidalysin are unable to damage epithelial cells or activate key signalling mechanisms that result in alarmin release and inflammatory responses and the recruitment of neutrophils ^22–24^. Consequently, neutrophil recruitment is severely impaired in models of mucosal and systemic candidiasis in response to candidalysin-deficient mutant strains ^25–28^. Thus, we investigated whether candidalysin can directly act on neutrophils and whether the toxin shapes neutrophil responses, which in turn may impact the outcome of invasive candidiasis.

We found that candidalysin-expressing *C. albicans* strains induce more NETs than candidalysin-deficient strains, indicating that candidalysin contributes to NET formation. Notably, synthetic candidalysin induces leukotoxic hypercitrullination and the release of NLS. NLS were dependent on NADPH oxidase-mediated reactive oxygen species (ROS) production and PAD4-mediated histone citrullination, but candidalysin did not induce cell cycle activation as indicated by lack of lamin A/C phosphorylation. Our data reveal that candidalysin is a critical virulence factor shaping neutrophil responses, which are essential for antifungal immunity.

## Results

### Candidalysin contributes to *C. albicans* induced NET formation

Neutrophils release NETs as a defense mechanism in response to *C. albicans* infections, particularly to control filamentous hyphae that are difficult to phagocytose ^2,5,29^. To investigate the impact of candidalysin on the neutrophil immune response towards *C. albicans*, we infected neutrophils with wild-type *C. albicans*, *ECE1*-deficient (*ece1*ΔΔ), and corresponding revertant (*ece1*ΔΔ+*ECE1*) strains, and a strain only lacking the candidalysin-coding sequence (P3) within the *ECE1* gene (*ece1*ΔΔ+*ECE1*-P3). After 4 h of infection, samples were prepared for indirect immunofluorescence microscopy to visualize extracellular trap events using decondensed neutrophil chromatin (DNA and α-histone) as marker (Fig. 1). Whereas wild-type and the revertant strain induced comparable amounts of NETs, the *ECE1*- and candidalysin-deficient strains triggered reduced levels (Fig. 1a). Based on previously published image-based quantitative analysis of NET formation ^30,31^, each DAPI-stained event exceeding 100 μm^2^ was considered a NET. The quantification revealed that both toxin-deleted strains induced significantly less NETs compared to toxin-expressing strains, with ~60% decreased levels after 3 and 5 h compared with the wild type (Fig. 1b). Notably, the scrutinized image-based quantification excluded background noise potentially derived from cell debris as confirmed by unstimulated control samples which were incubated in the same manner as stimulated samples (Fig. 1b). In conclusion, the data demonstrates that candidalysin contributes to NET formation triggered by *C. albicans* hyphae.

**Fig. 1.**
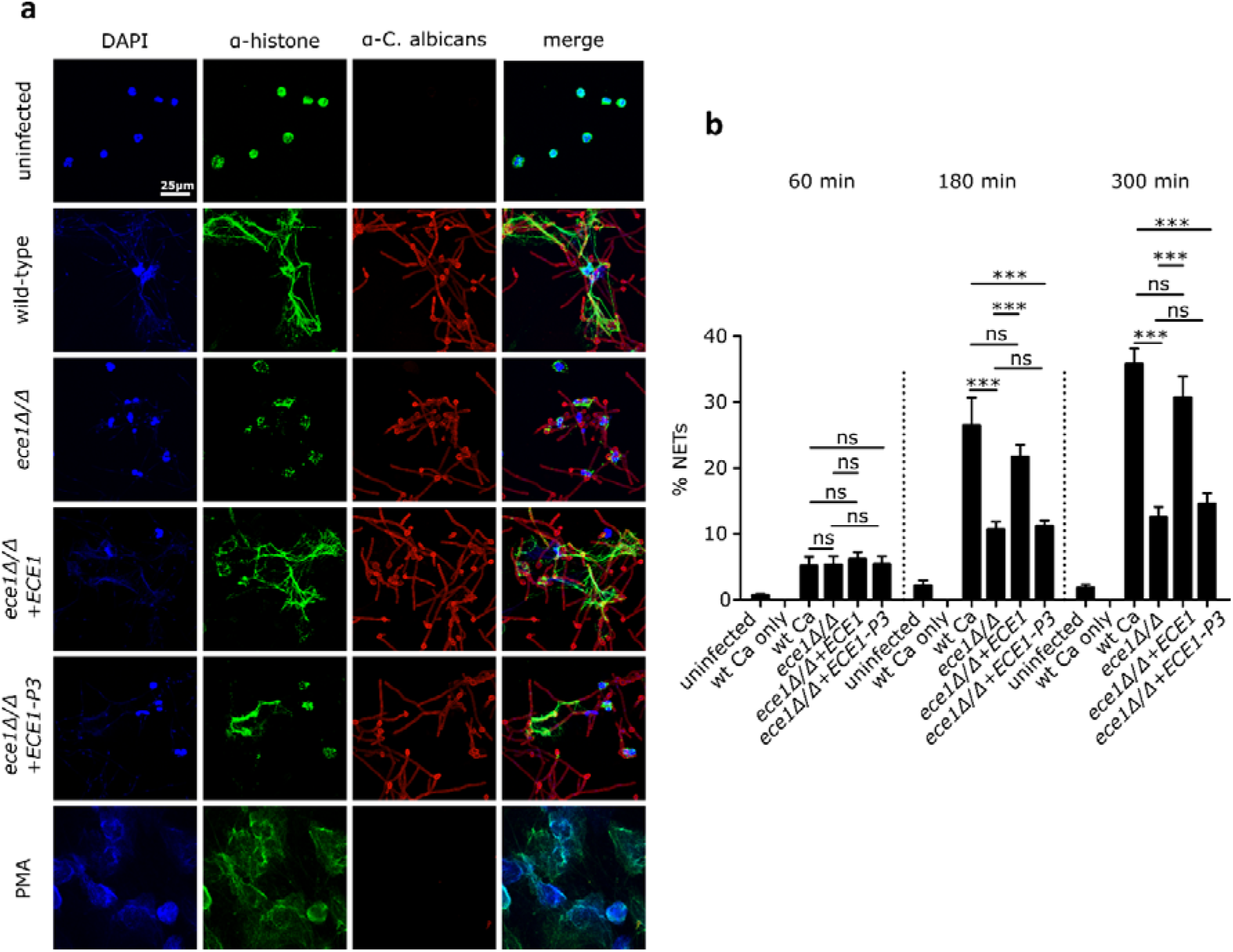
Candidalysin promotes NET formation. (a) Representative microscopic images (60X) of indirect immunofluorescence of human neutrophils 4 h after infection with wild-type and candidalysin deleted *C. albicans* strains (*ece1*Δ/Δ and *ece1*Δ/Δ+*ECE1*-P3). Lack of Ece1p/candidalysin production led to reduced NET formation as visualised by chromatin staining. Visual impression was corroborated with (b) quantitative image analysis of a time series experiment using ImageJ (n = 4, mean ± SEM). Each DAPI-stained event exceeding 100 μm^2^ was considered a NET. Statistical analysis conducted with two-way ANOVA with Bonferroni post-hoc test. Microscopic images are not obtained from the same experiment conducted for quantification due to different immunostaining procedures.

### Synthetic candidalysin induces NET-like structures

As candidalysin-expressing *C. albicans* strains induced more NETs, we aimed to investigate the potential of the toxin alone to stimulate neutrophils. Exposure of neutrophils to synthetic candidalysin was sufficient to trigger morphological changes (chromatin decondensation) in 46.3 ± 0.8% of cells after 4 h compared to 80.7 ± 3.2% after exposure to PMA, a well-known inducer of NETs (Fig. 2a). Neither scrambled candidalysin nor Ece1p peptide 2 (one of eight different Ece1p-derived peptides) affected neutrophil morphology, confirming specificity to candidalysin. Notably, the outspread structures in response to candidalysin were more compact, less fibrous and patchier compared to canonical NETs released upon stimulation with PMA or *C. albicans* hyphae (compare Fig. 2d with c and Figure 1a wild type, respectively). Hence, we concluded that synthetic candidalysin does not stimulate canonical NETs, but rather more compact DNA structures, resembling NLS that may be the result of leukotoxic hypercitrullination ^12,32^. Moreover, candidalysin demonstrated a dose-dependent effect with increased NLS formation from 3 μM to 15 μM. However, reduced NLS formation was observed at 70 μM (Fig. 2b), which can be explained by neutrophil cell death induced by the toxin as determined by a DNA Sytox Green assay (Fig. S1a). The structures induced by synthetic candidalysin were morphologically different from canonical NETs. However, the time course of morphological changes occurring during exposure to candidalysin was similar to the dynamics of morphological alterations during PMA-induced or *C. albicans* hypha-induced NET formation (Fig. S1b and Fig. 1a). In both cases, nuclear decondensation commenced at ~60 min and mixing of granular and nuclear components at ~120 min after stimulation (Fig. 2d and Fig. S2). In summary, synthetic candidalysin triggers morphologically distinct NLS in a time- and dose-dependent manner, whereas candidalysin-producing *C. albicans* hyphae induce canonical NETs (Fig. 1a).

**Fig. 2.**
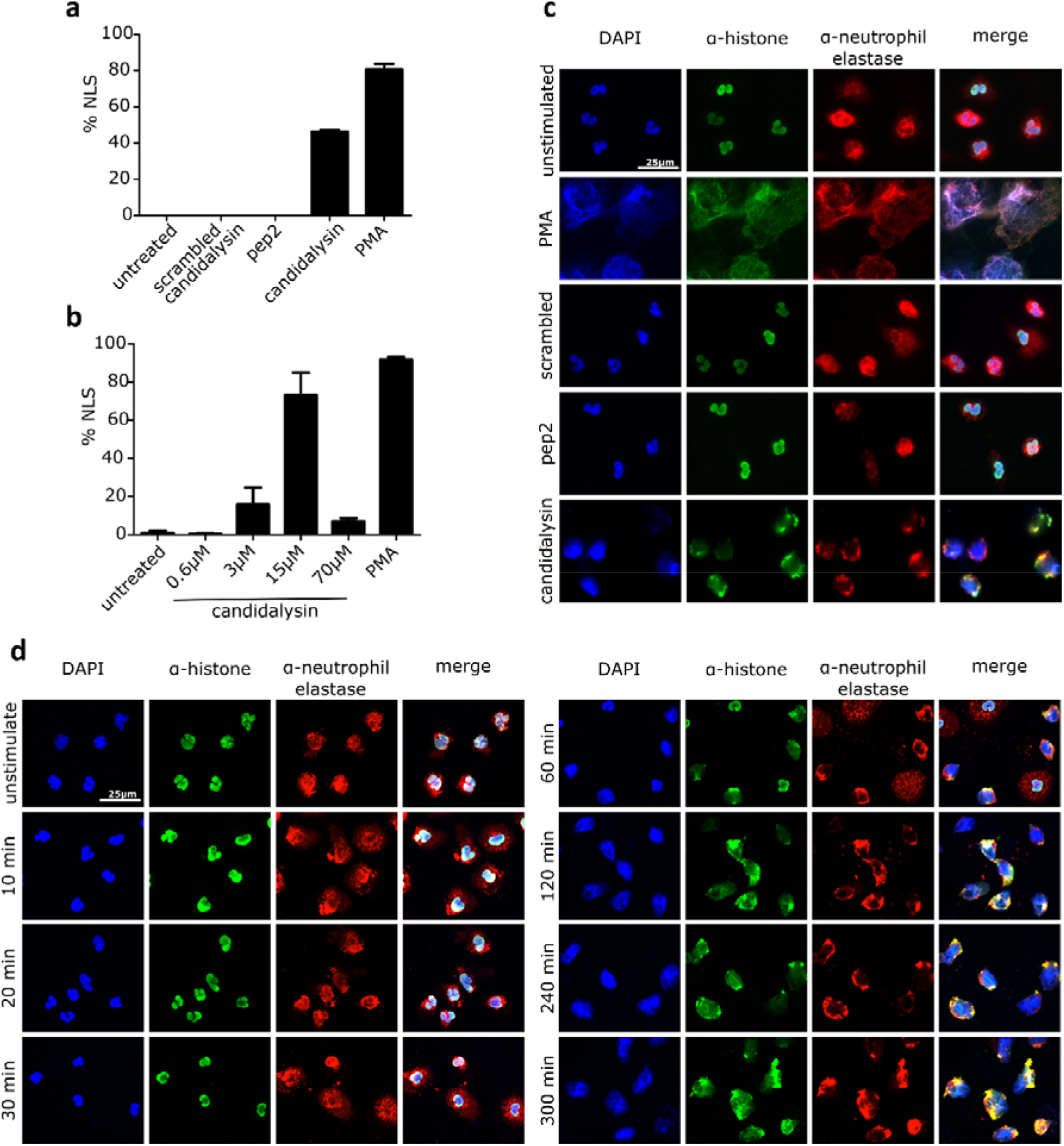
Synthetic candidalysin induces NLS in human neutrophils. Candidalysin, but not scrambled candidalysin or pep2, another Ece1p-derived peptide (all 15 μM), induce (a) DNA decondensation in human neutrophils after 4 h (n = 4) in a (b) dose-dependent manner (n = 3). NLS were quantified with the same criteria as previous described for NETs. Data shown as mean ± SEM. Confocal images (c) of immunostained cells display morphological changes involving nuclear and granular proteins after 4 h compared to unstimulated cells, or cells exposed to scrambled candidalysin and pep2. Time-dependent progression of morphological changes (d) in neutrophils induced by candidalysin over the course of 5 h (all images are with 60X magnification).

### Candidalysin-induced NET-like structures differ morphologically from NETs induced by various stimuli

To investigate candidalysin-triggered NLS further, we used scanning electron microscopy (SEM) that allows a more detailed view of the neutrophil-derived structures (**Error! Reference source not found**.a). To categorize the morphological alterations upon candidalysin stimulation, we compared the alterations with canonical ROS-dependent NETs triggered by PMA and NLS upon exposure to the bacterial peptide toxin ionomycin. Ionomycin has been previously reported to induce NLS, also referred to as leukotoxic hypercitrullination ^12,32^. Both, PMA and ionomycin generated widespread chromatin fibers in the extracellular space (Fig. 3a, left and middle panels). In contrast, fibrous, web-like structures were absent in candidalysin-treated neutrophil samples (Fig. 3a right panels, for 7 h treatment see Fig. S2). In addition, *C. albicans* hyphae induced NETs with observable fibers and threads similar to PMA- and ionomycin-stimulated neutrophils (Fig. 3b).

**Fig. 3.**
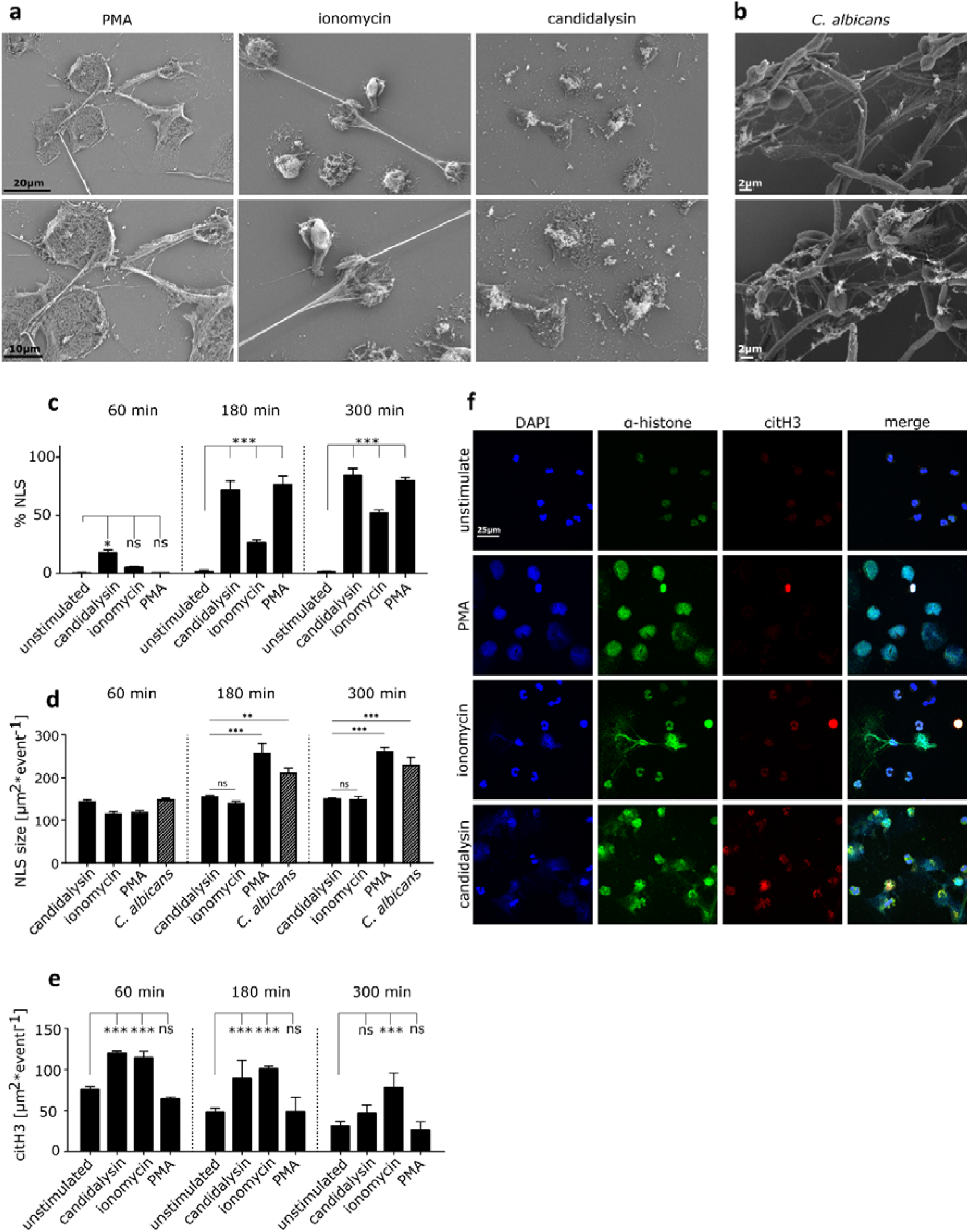
Morphological alterations triggered by candidalysin. (a) Scanning electron microscope images of candidalysin and ionomycin stimulated neutrophils after 3 h show differences in structural alterations compared to PMA-induced canonical NETs and (b) NETs induced by *C. albicans* hyphae (magnification (a) 3.00 KX on top, 5.00 KX at bottom and (b) 4.00 KX top, 3.00 KX bottom). Microscopic images were analysed by (c) amount of NLS formation (DNA decondensation), (d) average NLS size (only DNA-stained area > 100 μm^2^ considered) and (e) average histone citrullination level per event (n = 3, *C. albicans* n = 4). Data shown as mean ± SEM and statistical analysis conducted with two-way ANOVA with Bonferroni post-hoc test. (f) Representative immunofluorescence images 3 h after neutrophil stimulation support visually the quantitative data (60X magnification).

Image-based quantification of NLS events (candidalysin and ionomycin) and NETs (PMA and *C. albicans* hyphae) revealed that although candidalysin-triggered NLS appeared slightly earlier (after 1 h 17.9 ± 2.6% NLS), time dependency and quantity was similar compared to PMA-induced NETs (Fig. 3c). Ionomycin-induced changes, however, were more delayed with 26.5 ± 2.6% and 51.9 ± 3.1% NLS after 3 h and 5 h, respectively, and led to overall fewer NLS events. This was confirmed by an area-based analysis of the events (Fig. 3d). The average area per event exceeding 100 μm^2^ was determined using the images from the DNA stain. The frequency of extended threads was low for ionomycin-treated samples and the average area was significantly smaller for ionomycin-induced NLS (149.3 ± 6.21 after 3 h) than it was for PMA-induced (262 ± 8.43 after 3 h) and *C. albicans* hyphae-triggered NETs (231.34 ± 16.68 after 3 h). Lacking any recognizable fibers and threads candidalysin-triggered NLS displayed a lower average area per event (151.53 ± 0.62 after 3 h) comparable to ionomycin-triggered NLS (Fig. 3d).

The post-translational protein modification (PTM) of histones, in which arginine residues are enzymatically converted into peptidylcitrulline, was analysed since PTM is a driver of chromatin decondensation ^12^. The process of the PTM is called deamination or citrullination. Calcium influx activates protein arginine deiminase 4 (PAD4) and the enzyme subsequently facilitates histone citrullination (citH), which contributes to chromatin decondensation and eventually chromatin release. PAD4 activation has been reported for ionomycin ^32^ and nicotine ^31^. Thus, we assessed whether candidalysin induced histone citrullination in neutrophils. Indeed, like ionomycin, candidalysin increased histone citrullination in neutrophils above basal levels (**Error! Reference source not found**.e). Image quantification of histone citrullination using an antibody directed against citrullinated histone H3 (citH3), demonstrated that citH3 in candidalysin-stimulated neutrophils appeared more distributed than ionomycin-stimulated neutrophils, which remained concentrated in compact nuclei (Fig. 3f). Notably, we observed ~1.5-fold increased citH3 levels with ionomycin and candidalysin compared to unstimulated neutrophils and no increased citH3 levels with PMA (Fig. 3e). While citrullination levels decreased over time, ionomycin sustained high levels over 5 h. These data strongly suggest that candidalysin induces histone hypercitrullination in neutrophils, which likely promotes chromatin de-condensation.

### Candidalysin-expressing strains induce more NETs and higher citrullination levels than candidalysin-deficient strains

Since candidalysin contributed to the ability of *C. albicans* to induce NETs (Fig. 1) and synthetic candidalysin strongly stimulated histone citrullination, we investigated citrullination events when neutrophils were exposed to candidalysin-producing and candidalysin-deficient *C. albicans* strains. To assess this, neutrophils were stained with citrullination-specific antibodies (**Error! Reference source not found**.). Candidalysin-producing strains induced far more NETs than candidalysin-deficient strains (Fig. 4a). Image-based quantification corroborated the visual analysis and confirmed that candidalysin-producing *C. albicans* hyphae promote histone citrullination in neutrophils (Fig. 4b). As synthetic candidalysin only induces NLS, we concluded that candidalysin augments NET release when the toxin is secreted by *C. albicans* hyphae. Thus, we propose that the combination of candidalysin activity and fungal recognition via pattern recognition receptors ^33^ is required to fully trigger NET formation when neutrophils are exposed to *C. albicans in vivo*.

**Fig. 4.**
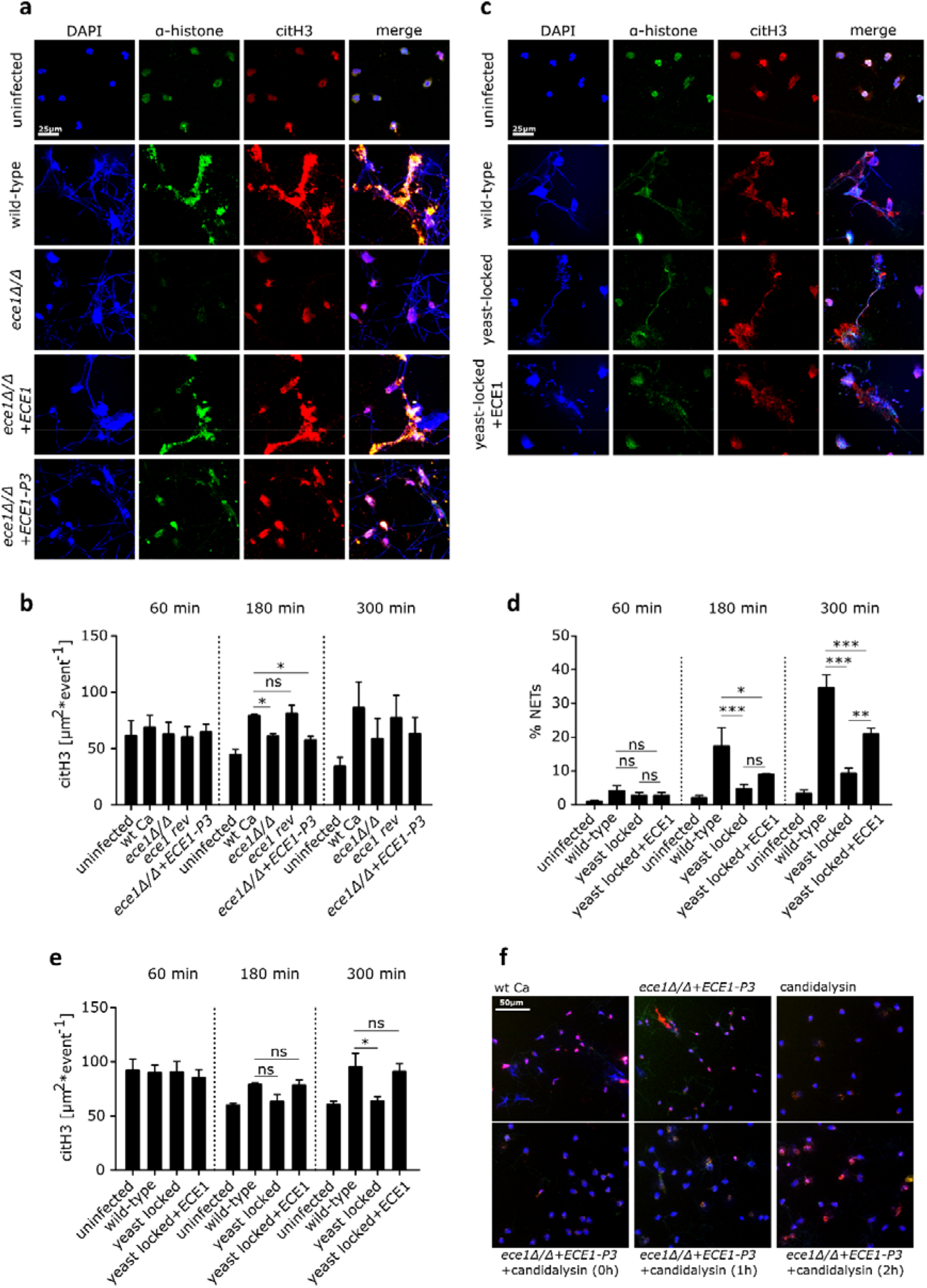
Candidalysin enhances NET formation through histone citrullination. (a) Representative immuno-fluorescence images (60X) of neutrophils infected with *C. albicans* wild-type and mutant strains after 3 h identified candidalysin as a major inducer of histone citrullination in human neutrophils with (b) significant decreased levels of citH3 in candidalysin-deficient strains (n = 4, mean ± SEM, statistical analysis with one-way ANOVA with Dunnett’s multiple comparison post-hoc test). (c, d) Although the yeast-locked mutant stimulated fewer NETs, *ECE1* overexpression partially recovered the potency (demonstrated by 40X microscopic images and image-based analysis) along with (e) increased histone citrullination. (d, e) Data of 4 donors shown as mean ± SEM and statistically analyses with one-way ANOVA with Bonferroni post-hoc test. (f) External addition of synthetic candidalysin resulted in a shift to NLS structures rather than NETs as visualized by microscopy after 5 h incubation (20X).

To test this proposition, neutrophils were infected with a yeast-locked (*cph1*ΔΔ /*efg1*ΔΔ) strain and an *ECE1*-overexpressing strain of the same genetic background *cph1*ΔΔ/*efg1*ΔΔ-*ECE1*) (Fig. 4c). As expected, the yeast-locked mutant induced significantly fewer NETs than wild-type *C. albicans* hyphae. Notably, the *ECE1*-overexpressing yeast-locked mutant, was partly restored in its ability to induce NET release, with 2-fold increased levels after 5 h compared to the yeast-locked mutant and over 60% of WT strain (Fig. 4d). This confirmed that candidalysin promotes *C. albicans*-triggered NET release. This was further confirmed by elevated citrullination patterns in presence of candidalysin (Fig. 4e). However, *ECE1*-overexpressing yeast-locked mutants were delayed in their ability to induce NET release and citrullination, which only emerged after 5 h of stimulation. Finally, we aimed to determine whether synthetic candidalysin could rescue NET formation when neutrophils were infected with a candidalysin-deficient strain (Fig. 4f). Interestingly, the addition of synthetic candidalysin resulted in a shift to NLS, irrespective of the time of addition, 1 h or 2 h post infection. The data suggest that candidalysin is the key driver of histone citrullination in neutrophils infected with *C. albicans*.

### NADPH oxidase enhances candidalysin-triggered NLS formation

As other peptide toxins can induce NETs independent of NADPH oxidase ^34^, we investigated the role for ROS in the induction of NLS by candidalysin. Treatment of neutrophils with synthetic candidalysin induced low levels of ROS, but significantly more than untreated neutrophils (Fig. 5a). In the strain context we observed lower ROS levels in the ece1Δ/Δ strain compared to its revertant strain, however, not to a significant extend (Fig. S3). Next, we assessed whether NADPH oxidase dependent ROS or mitochondrial ROS was induced by candidalysin using a luminol-based assay. Notably, candidalysin-induced ROS production was blocked by diphenyliodonium (DPI), a specific NADPH oxidase inhibitor, and by Tempol, a ROS scavenger. ROS inhibition was also observed with MitoTempo, a scavenger targeting mitochondrial ROS (Fig. 5b). The inhibitors alone had no significant effect on neutrophils (Fig. 5c). A similar pattern was observed in response to PMA (Fig. 5d). PMA activates protein kinase C (PKC) and the subsequent assembly and activation of NADPH oxidase ^35^. Thus, we concluded that candidalysin predominantly triggers NADPH oxidase to produce ROS but also moderate amounts of mitochondrial ROS. Next, we analysed how inhibition of ROS influenced the release of NLS triggered by candidalysin. Both, DPI and Tempol blocked candidalysin-induced ROS by 40-50%, while PMA-induced ROS production was totally blocked by DPI and Tempol (Fig. 5e). The data were confirmed by immunofluorescence where neutrophils were stained for DNA, histone, and elastase (Fig. 5f). Importantly, using NADPH oxidase-deficient neutrophils isolated from chronic granulomatous disease (CGD) patients (n = 3), we observed a reduction of candidalysin triggered NLS (30-40%) that was comparable to the effect of the ROS inhibitors (Fig. 5g and 5h). Together, the data confirm that candidalysin induces NLS in part in a NADPH oxidase-dependent fashion.

**Fig. 5.**
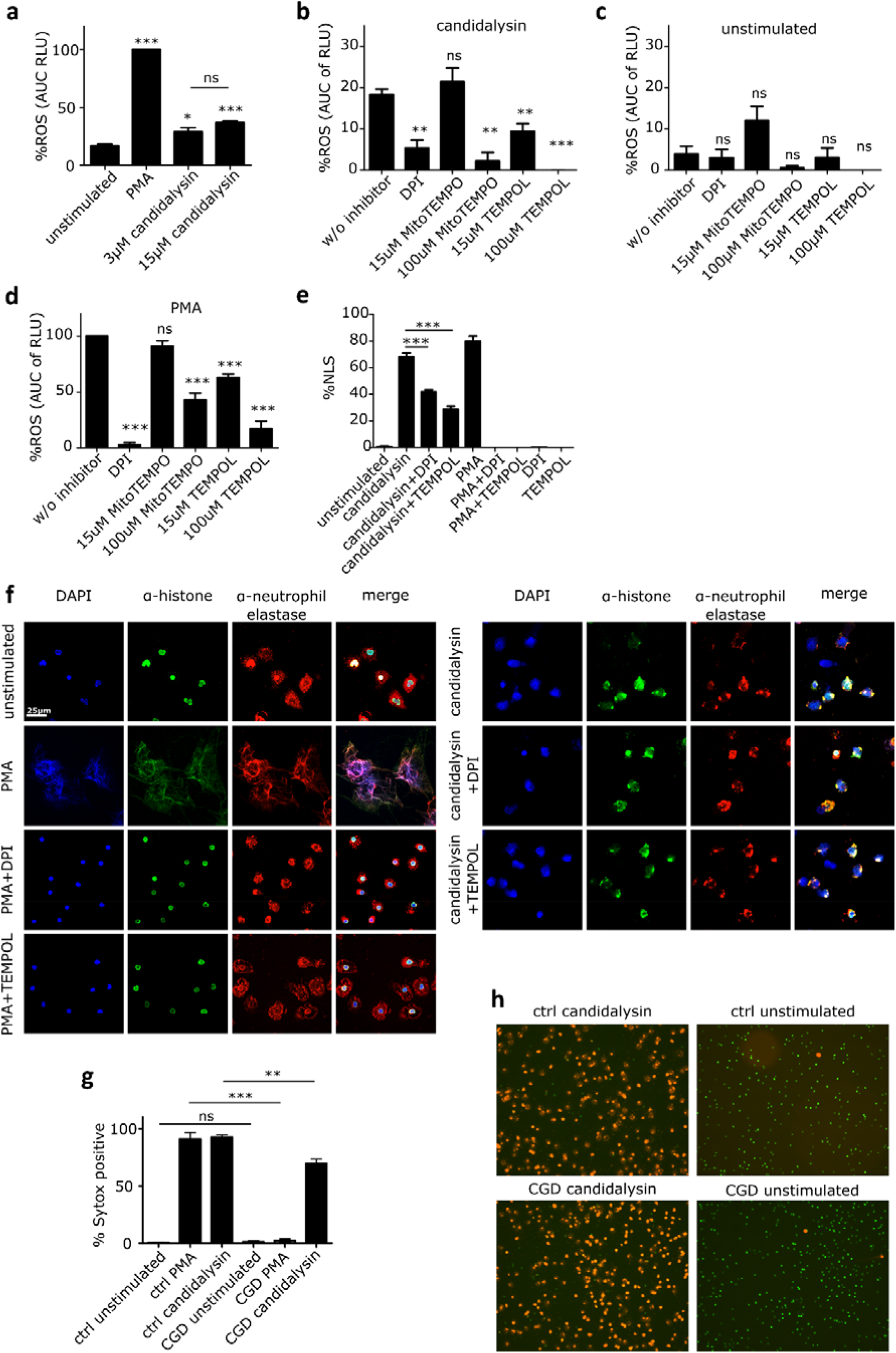
NLS induction by candidalysin is partially ROS-dependent. ROS response was measured in neutrophils upon stimulation with PMA and candidalysin (a) without and (b-d) in presence of a general ROS scavenger (TEMPOL), NADPH oxidase inhibitor (DPI) and a mitochondrial ROS inhibitor (MitoTEMPO) with a luminol-based assay. Data is presented as normalized area under the curve over 4 h treatment time (n = 3). The impact of stimulus-triggered ROS response on NLS formation was studied after 4.5 h incubation time with immunofluorescence microscopy with (e) image-based quantification (n = 3) and (f) a selection of representative images (60X magnification). (g) Sytox-positive cells after 4 h treatment. Candidalysin and PMA showed significantly decreased effects on neutrophils from CGD patients, as compared to neutrophils from healthy donors (n = 3). NLS responses were quantified using microscopic images of parallel staining using cell-impermeable Sytox Orange DNA dye (1 μM) to detect NETs/NLS and cell-permeable Sytox Green DNA dye (250 nM) to determine the total number of cells. (h) Representative images of the analysis are shown. Data shown as mean ± SEM and statistical analysis performed with One-way ANOVA with Bonferroni post-hoc test.

### Calcium influx and PAD4 activity contribute to candidalysin-triggered NLS

Cytoplasmic calcium (Ca^2+^) influx is required to stimulate PAD4 ^11^, which is responsible for histone citrullination and chromatin de-condensation during ionomycin-induced hypercitrullination. Since candidalysin also led to increased citrullination of histones in neutrophils, we aimed to elucidate the role of Ca^2+^ during candidalysin neutrophil interaction (Fig. 6). Candidalysin had a clear dose-dependent effect on intracellular Ca^2+^ influx (Fig. 6b). In contrast to Ca^2+^ spikes characteristic for chemokine receptor signalling, candidalysin-induced Ca^2+^ influx was not instantaneous (Fig. 6b and Suppl. Fig. S4) but started around 30 min post stimulation (Fig. 6b). This indicates that candidalysin most probably causes Ca^2+^ influx via pore formation and not via direct receptor stimulation. The PAD inhibitor Cl-amidine (PADi) reduced candidalysin-induced NLS formation by 70% after 180 min and by 50 % after 300 min, as quantified by microscopic analysis (Fig. 6c). Also, the cell-permeable calcium-chelator BAPTA-AM blocked candidalysin-induced NLS after 60 min (Fig. 6d). At later time points, BAPTA-AM led to an increase in NLS, probably due to toxic effects as indicated by higher background levels of NLS formation in non-stimulated, BAPTA-AM-treated neutrophils (Fig. 6d). We thus hypothesized that it may be possible to fully block candidalysin-induced NLS using a combination of PADi and the NADPH oxidase inhibitor DPI, since this combination would target both the ROS-dependent and -independent axis (Fig. 6a). At 180 min, the combination of PADi and DPI abrogated candidalysin-induced NLS slightly more than the individual inhibitors alone but not beyond the individual inhibitor effect at 300 min. Nevertheless, quantitative image analysis confirmed that PADi and DPI together blocked most NLS formation. For this purpose, neutrophils were stained with antibodies directed against histone H1, citrullinated histone H3, and with DNA dye DAPI. The analysis revealed that the treatment with DPI and PADi reduced the amount of patchy areas representing NLS after 300 min to almost background levels (Fig. 6e). Taken together, this suggests that candidalysin-induced NLS formation depends in part on both, ROS and PAD4.

**Fig. 6.**
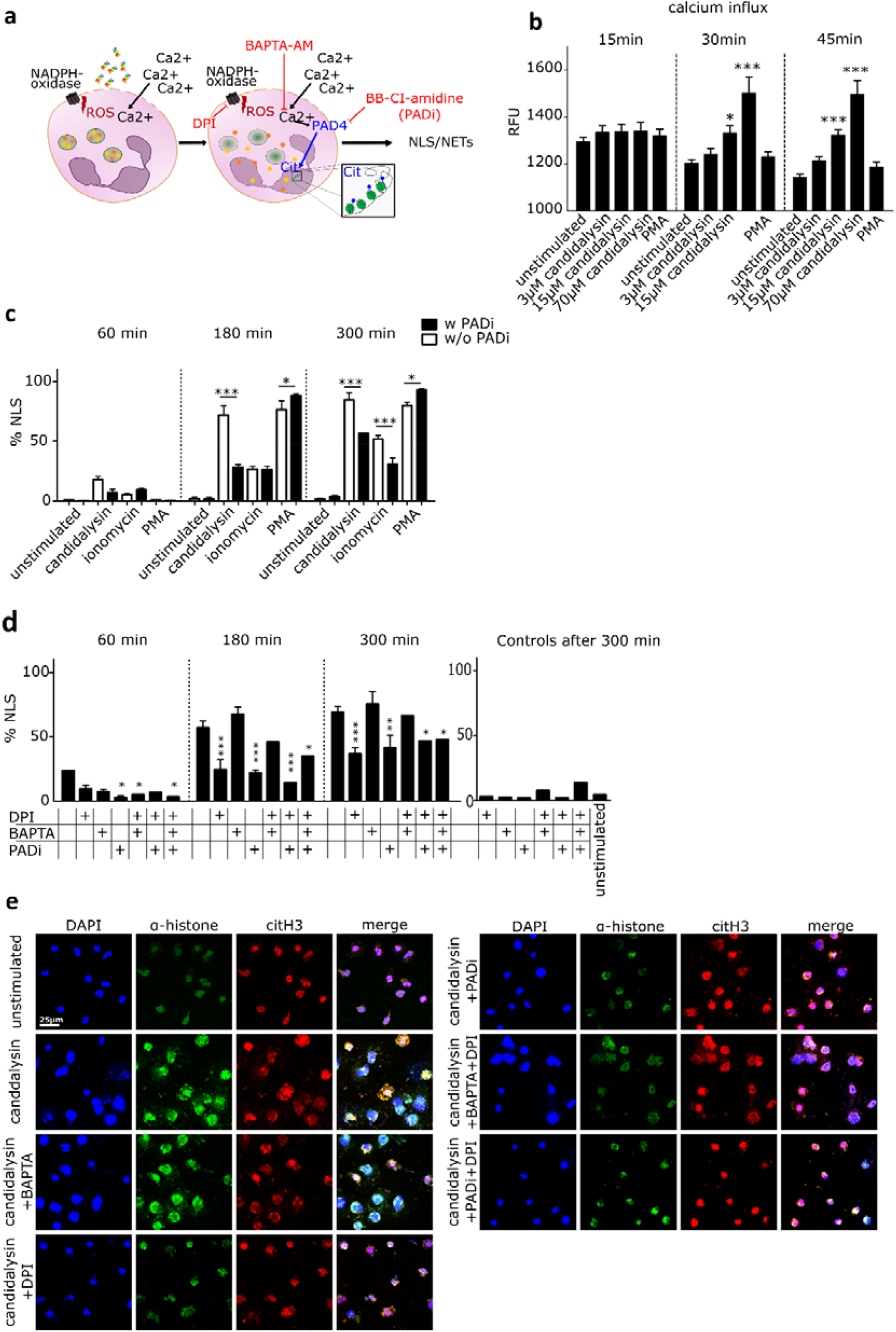
Candidalysin induces NLS via Ca^2+^- and ROS-dependent pathways. (a) Schematic image illustrating the suggested mechanisms by which candidalysin induces NLS in neutrophils. Both downstream effects of ROS and calcium-dependent PAD4 activation lead to chromatin decondensation. Inhibitors targeting NADPH oxidase (DPI) and PAD activation (BB-Cl-amidine, PADi) as well as calcium chelation (BAPTA) show effects. (b) Dose- and time-dependent calcium influx in neutrophils through candidalysin was measured with Fluo-8 AM (n = 4) and image-based quantification verified PAD-dependency of NLS formation via ionomycin and candidalysin (n = 3, data taken from same experiment as ***Error! Reference source not found.)***. (c-e) Combination treatment (DPI and PADi) blocking NADPH oxidase-dependent ROS and PAD-activation significantly reduced NLS formation through candidalysin (n = 3-4). Data shown as mean ± SEM and all statistical analysis performed with two-way ANOVA with Bonferroni post-hoc test. (e) Representative microscopic images (60X) demonstrate decreased morphological alterations through ROS and PAD blockage.

### Candidalysin initiates signalling pathways involved in NET formation

Our data show that candidalysin induced Ca^2+^ influx in neutrophils, which in turn activates PAD4 ^11,36^ leading to chromatin decondensation. Next, we investigated whether additional signalling pathways were involved in candidalysin induction of NLS (Fig. 7a). Phosphoinositide-3 kinase (PI3K) is a signalling molecule upstream of protein kinase B (Akt). PI3K and Akt are known molecular switches for neutrophil apoptosis or NET formation ^37^. The spleen tyrosine kinase (SYK), an important signalling protein involved in fungal detection, acts upstream of PI3K ^38^. In agreement, SYK signalling contributes to the regulation of NET formation triggered by *C. albicans* ^39^. As *C. albicans* hyphae bind to pathogen recognition receptors (PRRs), activate neutrophils and ultimately promote the release of NETs, we aimed to elucidate whether candidalysin alone leads to the activation of similar pathways in neutrophils. Hence, we stimulated neutrophils with candidalysin in the presence or absence of specific inhibitors for SYK, PI3K, and Akt. Interestingly, SYK blockade with R406 and PI3K blockade with wortmannin reduced NLS formation by candidalysin almost to background levels (Fig. 7b). The inhibitor piceatannol, which blocks both SYK and PI3K, also blocked NLS formation (Fig. 7d). In contrast, Akt blockade with AKT inhibitor XI only partially blocked candidalysin-induced NLS formation (Fig. 7c). This was expected, since Akt signals towards a ROS-dependent mechanism in neutrophils ^37^, which is not critical for NLS induction by candidalysin. In contrast, candidalysin stimulation of neutrophils induces Ca^2+^ influx, which leads to PAD4 activation (Fig. 6c). Candidalysin has been reported to induce inflammasome activation via NOD-like receptor family pyrin domain containing 3 (NLRP3) ^18^. However, NLRP3 activation seems to be dispensable for NLS induction (Fig. S5). Cell cycle molecules are also activated in the latter stages of NET formation and a hallmark of cell cycle induction is the phosphorylation of lamin A/C ^7^. However, unlike *C. albicans*, synthetic candidalysin did not trigger the phosphorylation of lamin A/C (Fig. 7e, f). Thus, we conclude that pathways involved in NET formation are triggered by candidalysin but these pathways cannot be fully sustained, thus NLS are formed rather than NETs. It is likely that a combination of candidalysin activity and hyphal recognition is required for sustained signalling, which will lead to complete chromatin decondensation and expulsion of NET fibers. This notion is confirmed by lack of downstream activation of the cell cycle proteins by synthetic candidalysin (Fig. 7c).

**Fig. 7.**
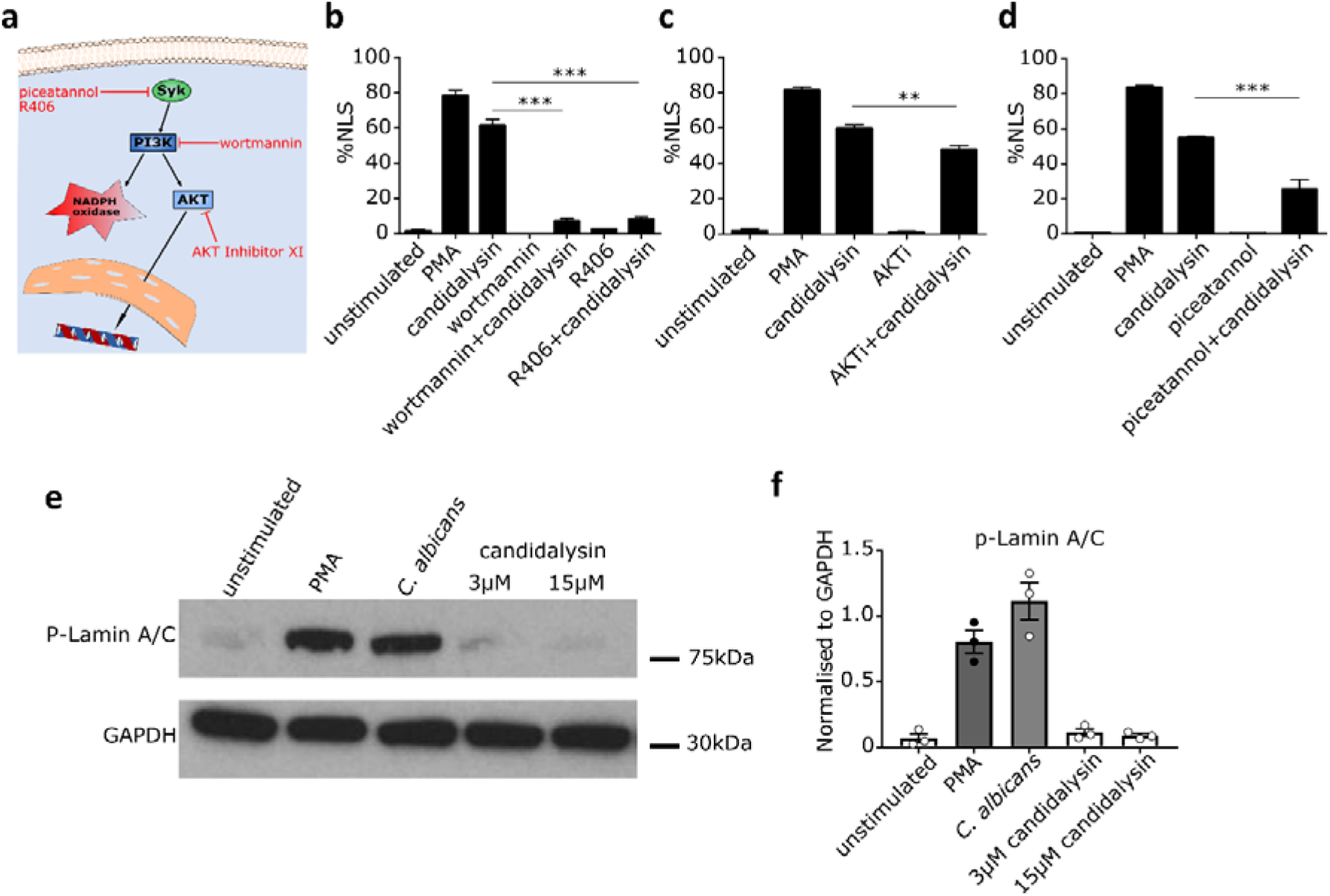
Candidalysin triggers signalling pathways involved in NET formation. (a) Schematic image shows the pathways involved in NET formation and inhibitors used to obtain mechanistic insights. (b-d) Blocking main kinases involved in NET formation with 15 μM R406 (SYK), 12.5 μM piceatannol (SYK), 15 μM wortmannin (PI3K) and 2.5 μM AKT inhibitor XI decreased NLS formation upon candidalysin stimulation in human neutrophils from healthy volunteers analysed using image analysis (n = 3, mean ± SEM, statistical analysed with one-way ANOVA with Bonferroni post-hoc test). (e) Western blot and (f) quantitative analysis (n = 3) did not show phospho-Lamin A/C activation by candidalysin.

### Neutrophils remain functional in the presence of candidalysin

Next, we investigated whether neutrophils exposed to candidalysin retain essential antimicrobial function, such as ROS production and phagocytosis. Although cellular death, as assessed using Sytox Green cell-impermeable DNA dye, occurs in increasing rates in candidalysin-treated neutrophils in a dose-dependent manner (Fig. S1a), neutrophils generally retained their functionality. Neutrophils were able to phagocytose beads in the presence of candidalysin (Fig. 8a), which was confirmed by time-lapse video (Movie S1), indicating that both Sytox-negative and Sytox-positive neutrophils remained functional. Candidalysin-treated neutrophils were tested for their capacity to mount ROS using PMA as a stimulant. Production of ROS was evident, even 1 or 2 h after candidalysin treatment (Fig. 8b). Untreated neutrophils, which were allowed to rest for the times indicated between 30 min and 3 h, showed increased ROS responses upon PMA stimulation (Fig. 8b). Notably, even after 2 h treatment with 15 μM candidalysin, neutrophils remained responsive, with ~40-50% of the ROS generated by PMA-stimulated neutrophils in the absence of candidalysin. The data indicates that the majority of neutrophils do not die upon exposure to 15 μM candidalysin. Finally, ionomycin-treated cells showed a minor ROS response and were subsequently unable to produce ROS in response to PMA (Fig. S6).

**Fig. 8.**
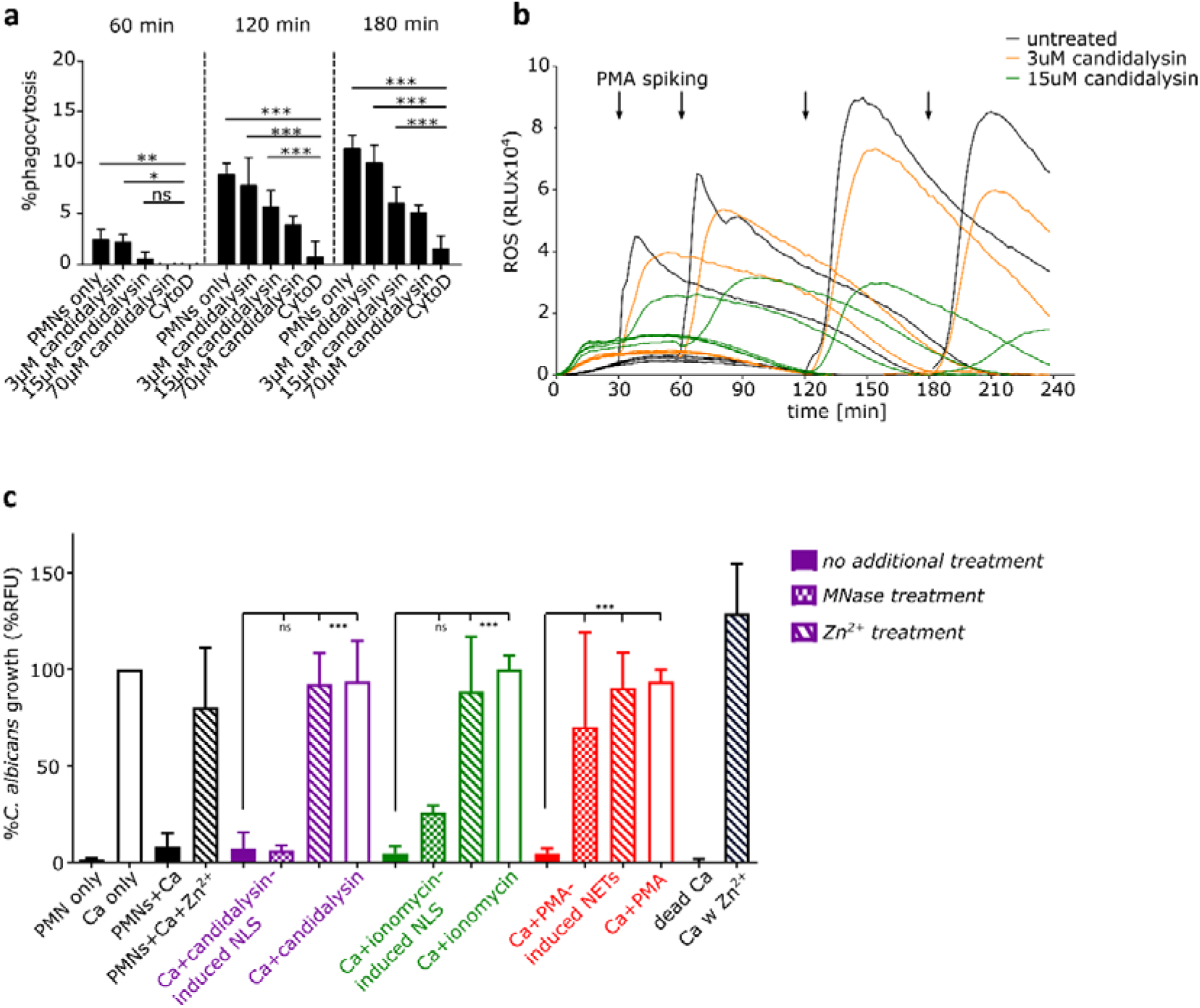
Candidalysin does not abrogate neutrophil functionality and NLS suppress fungal growth. (a) Despite cytotoxic effect of candidalysin on neutrophils, the immune cells were still able to phagocytose pre-opsonized zymosan-coated beads in presence of candidalysin, with significant higher levels compared to CytoD treated cells (one representative of 4 donors shown, statistical analysis performed with two-way ANOVA with Bonferroni post-hoc test). (b) The ability of ROS production in candidalysin-treated neutrophils was assessed over time through PMA spiking (one representative of 3 donors shown). (c) The antimicrobial activity assay revealed a similar fungal growth inhibition of NET-like structures induced by candidalysin and ionomycin as canonical PMA-NETs. *C. albicans* (Ca) growth on pre-induced NLS/NETs was measured with Calcofluor White staining after 16 h. The addition of Zn^2+^ to candidalysin-induced NLS before adding *C. albicans* negated the antimicrobial effect in opposite to no response to MNase exposure (n = 4, with following exception: n = 4 for MNase and Zn^2+^ treatment and only 2 donors for Zn^2+^ treatment on IOM-induced NLS).

### Candidalysin-triggered NLS inhibit *C. albicans* growth

As NETs inhibit the growth of *C. albicans*^29,40^, we investigated whether candidalysin-induced NLS harbored antifungal activity. Thus, we designed an image-based assay to assess *C. albicans* growth by quantifying calcofluor white staining in the presence of neutrophils that had been stimulated by candidalysin. Most importantly, the NLS triggered by candidalysin showed a strong anti-*Candida* effect (Fig. 8c). Optical density (OD) measurements were used to quantify biomass increase of *C. albicans* (Fig. S6). This confirmed that synthetic candidalysin did not suppress *C. albicans* growth, thus this effect was solely due to the candidalysin-induced NLS (Fig. S6). In addition, the *C. albicans* growth suppression could be reverted by addition of excess Zn^2+^ but not by micrococcal nuclease (MNase) (Fig. 8c). This confirms that, in contrast to canonical NETs, candidalysin-triggered NLS cannot be dismantled and removed by nuclease treatment, probably because NLS are considerably more compact than NETs. Therefore, NLS possessed antimicrobial effects even after nuclease treatment. The anti-*Candida* effect is most probably exerted via the zinc binding protein, calprotectin, as supplementation with excess Zn^2+^ blocked the antimicrobial effect of candidalysin-triggered NLS ^5^.

## Discussion

Candidalysin is the first fungal peptide toxin identified in any human fungal pathogen ^17^ and is critical for initiating inflammatory responses that trigger neutrophil recruitment during mucosal and systemic experimental candidiasis ^18,25–28^. As candidalysin is only produced by *C. albicans* hyphae ^41^, we investigated neutrophils response when these phagocytes encounter candidalysin-expressing *C. albicans* hyphae or synthetic candidalysin. Hyphae of candidalysin-expressing strains induced more NETs than *ECE1*-deficient and candidalysin-deficient strains (Fig. 1), indicating that candidalysin promotes NET formation. However, incubation of neutrophils with synthetic candidalysin was not sufficient to induce NETs (Fig. 2). Rather, stimulation with candidalysin led to citrullination of histones via PAD4, leukotoxic hypercitrullination, and the release of NLS. In contrast to canonical NETs, NLS are more compact and patchier with fewer clear fibers and threads (Fig. 3). Nevertheless, candidalysin-induced NLS did not occur instantaneously. Morphological changes were visible after 30-60 min exposure to candidalysin and intracellular mixing of granular and nuclear material was observed after ~120 min. After 300 min, ~80% of the neutrophil stimulated with candidalysin released NLS (Fig. 3). The role of candidalysin to promote canonical NET release was confirmed using a yeast-locked strain overexpressing *ECE1* (Fig. 4). While this overexpression construct did not reach the *ECE1* expression levels driven by the endogenous *ECE1* promoter, it nevertheless secreted significant level of candidalysin as described previously ^42^.

Interestingly, candidalysin induced low activity of NADPH oxidase and consequently ROS production. In CGD patient neutrophils, candidalysin-induced NLS were significantly reduced compared to control neutrophils; however, 60% of the CGD neutrophils were still able to release NLS (Fig. 5). Hence, while NADPH oxidase activity promotes candidalysin-induced NLS, it is not essential for NLS formation. This notion is clinically confirmed by the observation that CGD patients very rarely acquire *C. albicans* infections ^43^. In addition to ROS effects, candidalysin also induces the influx of calcium ions into the cytosol of neutrophils. Calcium influx is a known inducer of PAD4, the enzyme responsible for histone citrullination ^32^. We show that PAD4 is also required for histone decondensation during candidalysin-induced NLS formation (Fig. 6). It is unlikely that calcium influx in neutrophils is due to candidalysin directly triggering chemokine receptors, since calcium influx is slow and over time, and not in a pulse-like fashion.

High concentrations of candidalysin (70 μM) lyse human neutrophils more rapidly than lower concentrations (15 μM and 3 μM). Rapid lysis does not allow for regulated cellular processes to be induced within neutrophils. However, at lower concentrations (15 μM), the neutrophils encountering the toxin were still functional (ROS, phagocytosis) and mount a specific response leading to ROS production, PAD4 activation and the release of NLS. Signalling pathways involved in NET formation were also triggered by candidalysin (Fig. 7). Notably, SYK and PI3K inhibition significantly reduced the amount of candidalysin-triggered NLS; both signalling molecules are also inducers of NADPH oxidase ^37,39^. While chelation of calcium ions and PAD4 inhibition also reduced NLS formation, cell cycle processes such as the phosphorylation of lamin A/C, which is essential for the release of *C. albicans*-induced canonical NETs ^7^, were not activated by synthetic candidalysin. This indicates that candidalysin activates NET signalling pathways but these are not sustained or sufficient to induce the release of canonical NETs (Fig. 7). This notion is consistent with previous findings describing that PAD4 is dispensable for NET formation induced by *C. albicans* hyphae ^44^. Our data demonstrates that candidalysin is the main driver of histone citrullination in neutrophils infected with *C. albicans*. Lack of candidalysin production in *C. albicans* results in significantly reduced histone citrullination, accompanied with decreased NET formation. However, citrullination is not required for NET release, but rather governs the formation of NLS, which is dominant when candidalysin is added exogenously. With regard to *C. albicans* hyphae secreting candidalysin, it may be difficult to discriminate NLS form NETs, as both will be induced concurrently ^10^. It seems logical that the pore-forming activity of candidalysin augments the release of NET fibers during *C. albicans* infection, where PRRs will additionally be triggered on neutrophils, resulting in combinatorial activation of downstream pathways. In line with this notion, candidalysin drives histone citrullination, which contributes to chromatin decondensation. On the contrary, when neutrophils are exposed to synthetic candidalysin, the activation of pathways involved in NET formation are insufficiently sustained, resulting in the emergence of NLS. Importantly, our discovery that hyphae induce NETs and candidalysin induces NLS, provides an explanation for why opsonized *C. albicans* induce NETs in a ROS-dependent fashion, whereas un-opsonized *C. albicans* induce NETs in an ROS-independent fashion ^45^. As such, it appears that candidalysin has a more dominant effect in experimental settings without serum opsonization and a less dominant effect in the presence of serum opsonization.

It is noteworthy that candidalysin-induced NLS displayed anti-*Candida* activity. While some reports describe NLS as having no antimicrobial activity ^10^, we clearly see an anti-*Candida* effect by candidalysin-triggered NLS. As epithelial cells are able to expunge candidalysin for protection while *C. albicans* hyphae remain adeherent ^46^, recruited neutrophils may encounter candidalysin before direct contact with hyphae. In addition, neutrophil recruitment is virtually absent in mucosal and systemic models of candidasis in response to candidalysin-deficient strains ^25–28^. Hence, we chose to delineate the capacity of candidalysin-exposed neutrophils to kill *C. albicans*. Interestingly, candidalysin-triggered NLS are resistant to nuclease treatment but the resulting anti-*Candida* effect was Zn^2+^-dependent, indicating that growth inhibition of *C. albicans* by NLS relies on the presence of S100A8/A9 (calprotectin) ^5^ and potentially other Zn^2+^-binding neutrophil proteins. Given that NLS are morphologically distinct from NETs, being more compact and lacking threads, this likely explains why NLS are more resistant to nucleases. In context of *C. albicans* infection, candidalysin-induced permeabilization of the plasma membrane will result is large amounts of S100A8/A9 to be released, which will entangle in the structures. Further studies will be required to elucidate the key factors contributing to the anti-*Candida* effect of candidalysin-induced NLS.

In summary, this study shows that candidalysin promotes NET formation during *C. albicans* infection and NLS formation when present alone. During *C. albicans* infection, candidalysin drives the release of extracellular chromatin structures in both ROS-dependent and ROS-independent mechanisms, providing a possible rationale for the virtual absence of severe *C. albicans* infection in CGD patients. Importantly, neutrophils remain functional in the presence of candidalysin as both NETs and NLS display anti-*Candida* activity. Our findings serve as good starting point to further unravel the complexity of NET induction triggered by *C. albicans* and indicate that a combination of candidalysin(this study)and hyphal recognition ^2,3,33,47^ drive NET formation during *C. albicans* infection.

## Methods

### Fungal strain culture

The *Candida albicans* strains used in this study are listed in Table 1. In all cases, *C. albicans* was incubated in synthetic complete dropout medium (SC medium) for 16 h at 30°C. If not otherwise stated, a fresh subculture was inoculated in SC medium for 3 h before finally being washed three times with PBS, counted and adjusted according to each experimental protocol.

**Table 1.** Overview of *Candida albicans* strains used in this study. Fungal strains are described with parental cell line and genetic background.

| Fungal strain | Parental strain | Description | Genotype | Reference |
| --- | --- | --- | --- | --- |
| SC5314 |  | *Candida albicans* standard wild-type strain |  | <sup>48</sup> |
| *BWP17+Clp30* |  | Wild-type strain | ura3::λimm434/ura3::λimm434<br>arg4::hisG/arg4::hisG<br>his1::hisG/his1::hisG + Cip30 | <sup>49</sup> |
| *ece1ΔΔ* | *BWP17+Cip30* | *ECE1* knockout | *ece1::HIS1/ece1::ARG4*<br>*RPS1/rps1::URA3* | <sup>17</sup> |
| *ece1ΔΔ + ECE1* | *BWP17+Cip30* | *ece1Δ* revertant | *ece1::HIS1/ece1::ARG4*<br>*RPS1/rps1::(URA3 ECE1)* | <sup>17</sup> |
| *ece1ΔΔ + ECE1ΔIII* | *BWP17+Cip30* | *C. albicans*<br>*BWP17-Clp30*<br>with candidalysin<br>knockout | *ece1::HIS1/ece1::ARG4*<br>*RPS1/rps1::(URA3 ECE1<sub>Δ184-279</sub>)* | <sup>17</sup> |
| *cph1ΔΔ*<br>*/efg1ΔΔ* | *CAI4+Cip10* | *C. albicans* yeast-locked mutant<br>strain | *cph1::hisG/cph1::hisG*<br>*efg1::hisG/efg1::hisG-URA3-*<br>*hisG* | <sup>50</sup> |
| *cph1/efg1*<br>*ECE1*<br>Overexpression | *CAI4+Cip10* | *C. albicans* yeast-locked mutant<br>strain with *ECE1*<br>overexpression | *cph1/efg1 pENO1_ECE1* | <sup>42,51</sup> |

### Isolation of human polymorphonuclear neutrophils (PMNs)

Blood sampling for research purposes was conducted in accordance with the principles stated in the Declaration of Helsinki, and with agreement with the blood-central of the University Hospital of Umeå. CGD patient blood collection was approved by the ethical committee of Charité University Hospital, Berlin, Germany. Venous blood samples were drawn from healthy volunteers and CGD patients in EDTA tubes and neutrophils isolated as previously described ^52^. In brief, the neutrophil fraction was obtained using density centrifugation in Histopaque 1119 (Sigma-Aldrich) to separate granulocytes followed by a discontinuous Percoll (GE Healthcare Life Sciences) gradient to isolate neutrophils. After RBC lysis (RBC lysis buffer, BioLegend) the cells were resuspended in RPMI 1640 media (Lonza, supplemented with 5% HEPES) and counted. Only neutrophils with the viability above 90% were selected for further experimentation.

### Neutrophil stimulation

Neutrophils were seeded on glass cover slips coated with 0.001% poly-L-lysine (Sigma-Aldrich) with a concentration of 1 × 10^5^ cells per well if not stated otherwise. PMNs were stimulated with 4 μM ionomycin (free acid, Sigma-Aldrich), 100 nM Phorbol 12-myristate 13-acetate (PMA, Sigma-Aldrich), 15 μM Ece1 peptides including candidalysin (if not otherwise stated) or infected with *C. albicans* yeast (MOI 2) for a defined time period, following fixed using 2% paraformaldehyde (PFA) and stored at 4°C. In the infection experiments, the fungus was added to 1 × 10^4^ PMNs 1 h after the cells were seeded.

For the pathway studies neutrophils were incubated for 30 min before stimulation with 10 μM BB-Cl-amidine (PADi, Cayman Chemicals), 15 μM Dipenyleneiodium (DPI, Sigma-Aldrich), 15 μM 4-Hydroxy-TEMPO (TEMPOL, Sigma-Aldrich), 10/20 μM BAPTA-AM (Abcam), SYK inhibitors R406 (15 μM, InvivoGen) and piceatannol (12.5 μM, InvivoGen), 15 μM PI3K blocker wortmannin (InvivoGen), 2.5 μM AKT inhibitor XI (InvivoGen) or NLRP3 blockage using compound MCC950 (1 μM, InvivoGen).

### Immunostaining, Microscopy and Quantification

For immune staining the cover slips were washed with PBS, cells permeabilized with 0.5% TritonX-100 (company) for 1 min and then blocked at room temperature for 30 min in 3% bovine serum albumin (Sigma-Aldrich) buffer. Antibodies directed against histone H1 (final 1 μg/mL, #BM465, Acris) and citrullinated histone H3 (citrulline R2+R8+R17, 1 μg/mL, ab5103, Abcam) were applied and incubated for 1 h at 37°C following by secondary antibodies conjugated with Alexa Fluor dyes 488 and 568 (10 μg/mL, Thermo Fisher). DNA was stained with DAPI (1 μg/mL, Sigma-Aldrich). Prolong Diamond Antifade Mountant (Invitrogen) was used for mounting. For visualization and quantification 10 to 14 images per condition with around 50 to 150 cells were randomly taken with 20X magnification (Nikon Eclipse 90i fluorescence microscope with NIS Elements software) and the analysis performed with ImageJ.

For quantification of NETs and NET-like structures (NLS) (modified accordingly ^30,31^), DAPI stained events with an area over 15 μm^2^ were measured and nuclei exceeding 100 μm^2^ were counted. For quantification of citrullinated histone the Alexa Fluor 568 total stained area was measured and further normalized as signal per cell based on the event count of the DNA staining. NETs are characterized as web-like structures with threads spanning over several dozens of micro meter, whereas NLS are more compact, patchy and without longer threads.

Confocal images were taken with Nikon A1R confocal (LSM) controlled by Nikon NIS Elements interface with a Nikon Eclipse Ti-E inverted microscope using 60X magnification.

To quantify NLS from CGD patient neutrophils in comparison to neutrophils from healthy individualss, cells were seeded in a concentration of 1 × 10^5^cells per well in 24-well plates in RPMI medium. Neutrophils were stained using cell-impermeable Sytox Orange DNA dye (1 μM, Thermo Fisher) to detect NETs and cell-permeable DNA dye Sytox Green (250 nM, Thermo Fisher) to determine the total number of cells. NETs/NLS were imaged at 4 h post stimulation using 20X magnification on a EVOS FL Auto Microscope (Thermo Scientific).

### Scanning Electron Microscopy

Neutrophils were stimulated as described above. After fixation, the cells were washed with PBS and subsequently dehydrated in a series of graded ethanol (70, 80, 90, 95 and 100%). After critical point drying with Leica EM CPD300, the cover slips were coated with a 2 nm platinum layer (Quorum Q150T-ES Sputter Coater). Representative images were acquired using field-emission scanning electron microscopy (SEM, Carl Zeiss Merlin) with secondary electron detector at accelerating voltage of 4 kV, probe current of 120 pA and a working distance of 5.1 mm.

### Western Blot

2 × 10^6^ neutrophils were stimulated with 100 nM PMA, opsonised *C. albicans* (MOI 5) or candidalysin (3 μM and 15 μM) for 90 min in tubes, followed by centrifugation at 400 × g for 5 min. Neutrophils were then resuspend in 40 μl PBS supplemented with 1x Protease and phosphatase inhibitor (Thermo Fisher) and placed on ice for 10 min. Subsequently, SDS was added, samples were boiled at 100°C for 10 min, sonicated with 3 pulses of 15 s at 100% power (QSonica) and stored at −20°C. 10 μl of sample were loaded in a 4-12% Bis-Tris pre-cast gel (Invitrogen). Gel was transferred to a PVDF membrane and blocked in 1% BSA (Fisher) in TBST, followed by blotting with anti-phospho-lamin A/C (1:1000, Cell Signaling #13448) and anti-GAPDH (1:1000, Cell Signaling #2118).

### Cell Death Assay

Neutrophil cell death or the presence of extracellular DNA was quantified using a Sytox Green-based (Invitrogen) fluorescence assay similar to previous descriptions ^2,35^. Briefly, cells were seeded in a black 96 well plate with a concentration of 5 × 10^4^ cells per well. Subsequently, Sytox Green, a membrane-impermeable DNA dye, was added to a final concentration of 5 μM, before cells were stimulated. The fluorescence signal was measured in a plate-based fluorescence spectrophotometer (Fluostar Omega, BMG) at 37°C and 5% CO_2_ for 10 h in intervals of 10 min. The percentage of dead cells was calculated using TritonX-100 lysed neutrophils as 100% control. Each experiment was performed in 4 replicates.

### ROS measurement

The induction of ROS was measured by oxidation of luminol and determined in Varioskan Flash reader (Thermo Fisher Scientific) at 37°C. 5 × 10^4^ PMNs per well were seeded into black 96 well plates and incubated in media containing 50 mM luminol (Sigma-Aldrich), 1.2 U/well HRP (Sigma-Aldrich) and different inhibitors for 30 min at 37°C and 5% CO_2_. After stimulation or infection with *C. albicans* (MOI 2), the luminescence measurement was started and data was obtained every 2 min. Each experiment was performed in 4 replicates.

For ROS inhibition TEMPOL, MitoTEMPO and DPI (all from Sigma-Aldrich) were used at a concentration of 15 or 100 mM. For the functional assessment 100 nM PMA (Sigmal-Aldrich) was added to previously stimulated neutrophils after 30, 60 or 120 min.

### Phagocytosis Assay

Neutrophils (5 × 10^4^ cells/well) were seeded into a black 96 well plate and stimulated with different concentrations of candidalysin. After 30 min incubation time, 25 μg/well opsonized pHrodo Red Zymosan bioparticle conjugates for phagocytosis (Thermo Fisher) were added and the fluorescence intensity of the beads (excitation 560/emission 585 nm) was measured with Fluostar Omega plate reader (BMG). Acidized beads (phthalate buffer [100 mM; pH 4]) and PMNs with the blocked cytoskeleton (12.5 μM cytochalasin D) served as 100% and 0% control, respectively. Each experiment was performed in 4 replicates. Bead opsonization was performed with 60% human serum for 30 min and the control cells were incubated with CytoD for 80 min.

The time-lapse imaging (video attached) was performed with pHrodo Red *S. aureus* bioparticle conjugates for phagocytosis (Thermo Fisher) as described above in addition of final 5 μM Sytox Green. The video shows neutrophils 30 min after 15 μM candidalysin treatment.

### Antimicrobial Activity Assay

The growth inhibition of candidalysin pre-treated neutrophils on *C. albicans* was assessed with an end-point chitin staining with Calcofluor White (Sigma-Aldrich). Neutrophils (1 × 10^5^ cells/well) were seeded in a poly-L-lysine (Sigma-Aldrich) pre-coated 96 well plate and after 30 min incubation time stimulated with 15 μM candidalysin, 4 μM ionomycin or 100 nM PMA for 5 h. After treating designated wells with 10 U/mL MNase, the total well volume was removed and *C. albicans* in a concentration of 5 × 10^4^ cells/well (MOI 0.5) added. Thimerosal (Sigma-Aldrich)-killed Candida served as a control. Designated wells were supplemented with 5 μM ZnSO_4_ (Sigma-Aldrich) as a Zink source. The plate was incubated for 16 h at 37°C and 5% CO_2_, MNase added to wells previously not treated and subsequently the cells were fixed with 4% PFA for 20 min at room temperature. After Calcofluor White staining (0.1 mg/mL for 10 min), images were acquired with Cytation 5 Cell Imaging Reader (BioTek) and a cell number representative fluorescence signal obtained. Each experiment was performed in 4 replicates. To exclude an inhibitory effect of the toxins itself on *C. albicans*, wells were treated in absence of neutrophils and then infected with the fungus.

### Growth curve

To study the growth of *C. albicans* in presence of candidalysin, a measurement of optical density (λ=600) was performed. 15 μM candidalysin was added to a poly-L-lysine pre-coated 96 well plate as described, before being washed and infected with different concentrations of *C. albicans*, or directly added to the well together with the fungus. Data was obtained with Fluostar Omega plate reader (BMG) over 16 h with an interval of 1 h at 37°C and 5% CO_2_. Each experiment was performed in 4 replicates.

### Calcium influx

The measurement of calcium influx into cells was adapted from ^53^. After neutrophil isolation, the cells were resuspended in HBSS without Calcium and Magnesium (Lonza). 5 μM Fluo-8 AM (Abcam) was added to PMNs at 37°C for 90 min. Cells were washed once and resuspended in RPMI 1640. In a total reaction volume of 120 μl, 1 × 10^5^ cells were seeded into a black 96 well plate and stimulated with 70, 15, 3 and 0.56 μM candidalysin. After 10 min incubation, the fluorescence was measured (Ex490/Em520) for 60 min with Fluostar Omega plate reader (BMG). Each experiment was performed in triplicates.

### Statistical analysis

For all calculations and analyses GraphPad Prism Software 5.0 (GraphPad Software) was used. Bars represent 95% CI and p value significance is shown as following: *p < 0.05, **p < 0.01, ***p < 0.001. Numbers of biological replicates using independent neutrophil donors (n) are indicated in the figure label.

---

**Fig. 9. Candidalysin promotes NET formation**. (A) Representative microscopic images (60X) of indirect immunofluorescence of human neutrophils 4 h after infection with wild-type and candidalysin deleted *C. albicans* strains (*ece1*Δ/Δ and *ece1*Δ/Δ+*ECE1*-P3). Lack of Ece1p/candidalysin production led to reduced NET formation as visualised by chromatin staining. Visual impression was corroborated with (B) quantitative image analysis of a time series experiment using ImageJ (n = 4, mean ± SEM). Each DAPI-stained event exceeding 100 μm^2^ was considered a NET. Statistical analysis conducted with two-way ANOVA with Bonferroni post-hoc test. Microscopic images are not obtained from the same experiment conducted for quantification due to different immunostaining procedures.

**Fig. 10. Synthetic candidalysin induces NET-like structures in human neutrophils**. Candidalysin, but not scrambled candidalysin or pep2, another Ece1p-derived peptide (all 15 μM), induce (A) DNA decondensation in human neutrophils after 4 h (n = 4) in a (B) dose-dependent manner (n = 3). NLS were quantified with the same criteria as previous described for NETs. Data shown as mean ± SEM. Confocal images (C) of immunostained cells display morphological changes involving nuclear and granular proteins after 4 h compared to unstimulated cells, or cells exposed to scrambled candidalysin and pep2. Time-dependent progression of morphological changes (D) in neutrophils induced by candidalysin over the course of 5 h (all images are with 60X magnification).

**Fig. 11. Morphological alterations triggered by candidalysin**. (A) Scanning electron microscope images of candidalysin and ionomycin stimulated neutrophils after 3 h show differences in structural alterations compared to PMA-induced canonical NETs and (B) NETs induced by *C. albicans* hyphae (magnification (A) 3.00 KX on top, 5.00 KX at bottom and (B) 4.00 KX top, 3.00 KX bottom). Microscopic images were analysed by (C) amount of NLS formation (DNA decondensation), (D) average NLS size (only DNA-stained area > 100 μm^2^ considered) and (E) average histone citrullination level per event (n = 3, *C. albicans* n = 4). Data shown as mean ± SEM and statistical analysis conducted with two-way ANOVA with Bonferroni post-hoc test. (F) Representative immunofluorescence images 3 h after neutrophil stimulation support visually the quantitative data (60X magnification).

**Fig. 12. Candidalysin enhances NET formation through histone citrullination**. (A) Representative immuno-fluorescence images (60X) of neutrophils infected with *C. albicans* wild-type and mutant strains after 3 h identified candidalysin as a major inducer of histone citrullination in human neutrophils with (B) significant decreased levels of citH3 in candidalysin-deficient strains (n = 4, mean ± SEM, statistical analysis with one-way ANOVA with Dunnett’s multiple comparison post-hoc test). (C, D) Although the yeast-locked mutant stimulated fewer NETs, *ECE1* overexpression partially recovered the potency (demonstrated by 40X microscopic images and image-based analysis) along with (E) increased histone citrullination. (D, E) Data of 4 donors shown as mean ± SEM and statistically analyses with one-way ANOVA with Bonferroni post-hoc test. (F) External addition of synthetic candidalysin resulted in a shift to NLS structures rather than NETs as visualized by microscopy after 5 h incubation (20X).

**Fig. 13. NLS induction by candidalysin is partially ROS-dependent**. ROS response was measured in neutrophils upon stimulation with PMA and candidalysin (A) without and (B-D) in presence of a general ROS scavenger (TEMPOL), NADPH oxidase inhibitor (DPI) and a mitochondrial ROS inhibitor (MitoTEMPO) with a luminol-based assay. Data is presented as normalized area under the curve over 4 h treatment time (n = 3). The impact of stimulus-triggered ROS response on NLS formation was studied after 4.5 h incubation time with immunofluorescence microscopy with (E) image-based quantification (n = 3) and (F) a selection of representative images. (G) Sytox-positive cells after 4 h treatment. Candidalysin and PMA showed significantly decreased effects on neutrophils from CGD patients, as compared to neutrophils from healthy donors (n = 3). NLS responses were quantified using microscopic images of parallel staining using cell-impermeable Sytox Orange DNA dye (1 μM) to detect NETs/NLS and cell-permeable Sytox Green DNA dye (250 nM) to determine the total number of cells. (H) Representative images of the analysis are shown. Data shown as mean ± SEM and statistical analysis performed with One-way ANOVA with Bonferroni post-hoc test.

**Fig. 14. Candidalysin induces NLS via Ca^2+^ - and ROS-dependent pathways**. (A) Schematic image illustrating the suggested mechanisms by which candidalysin induces NLS in neutrophils. Both downstream effects of ROS and calcium-dependent PAD4 activation lead to chromatin decondensation. Inhibitors targeting NADPH oxidase (DPI) and PAD activation (BB-Cl-amidine, PADi) as well as calcium chelation (BAPTA) show effects. (B) Dose- and time-dependent calcium influx in neutrophils through candidalysin was measured with Fluo-8 AM (n = 4) and image-based quantification verified PAD-dependency of NLS formation via ionomycin and candidalysin (n = 3, data taken from same experiment as ***Error! Reference source not found***.C-E). Combination treatment (DPI and PADi) blocking NADPH oxidase-dependent ROS and PAD-activation significantly reduced NLS formation through candidalysin (n = 3-4). Data shown as mean ± SEM and all statistical analysis performed with two-way ANOVA with Bonferroni post-hoc test. (E) Representative microscopic images (60X) demonstrate decreased morphological alterations through ROS and PAD blockage.

**Fig. 15. Candidalysin triggers signalling pathways involved in NET formation**. (A) Schematic image shows the pathways involved in NET formation and inhibitors used to obtain mechanistic insights. (B-D) Blocking main kinases involved in NET formation with 15 μM R406 (SYK), 12.5 μM piceatannol (SYK), 15 μM wortmannin (PI3K) and 2.5 μM AKT inhibitor XI decreased NLS formation upon candidalysin stimulation in human neutrophils from healthy volunteers analysed using image analysis (n = 3, mean ± SEM, statistical analysed with one-way ANOVA with Bonferroni post-hoc test). (E) Western blot and (F) quantitative analysis (n = 3) did not show phospho-Lamin A/C activation by candidalysin.

**Fig. 16. Candidalysin does not abrogate neutrophil functionality and NLS suppress fungal growth.** (A) Despite cytotoxic effect of candidalysin on neutrophils, the immune cells were still able to phagocytose pre-opsonized zymosan-coated beads in presence of candidalysin, with significant higher levels compared to CytoD treated cells (one representative of 4 donors shown, statistical analysis performed with two-way ANOVA with Bonferroni post-hoc test). (B) The ability of ROS production in candidalysin-treated neutrophils was assessed over time through PMA spiking (one representative of 3 donors shown). (C) The antimicrobial activity assay revealed a similar fungal growth inhibition of NET-like structures induced by candidalysin and ionomycin as canonical PMA-NETs. *C. albicans* (Ca) growth on pre-induced NLS/NETs was measured with Calcofluor White staining after 16 h. The addition of Zn^2+^ to candidalysin-induced NLS before adding *C. albicans* negated the antimicrobial effect in opposite to no response to MNase exposure (n = 4, with following exception: n = 4 for MNase and Zn^2+^ treatment and only 2 donors for Zn^2+^ treatment on IOM-induced NLS).

