## Supplementary information for "*Candida albicans* promotes neutrophil extracellular trap formation and leukotoxic hypercitrullination via the peptide toxin candidalysin"

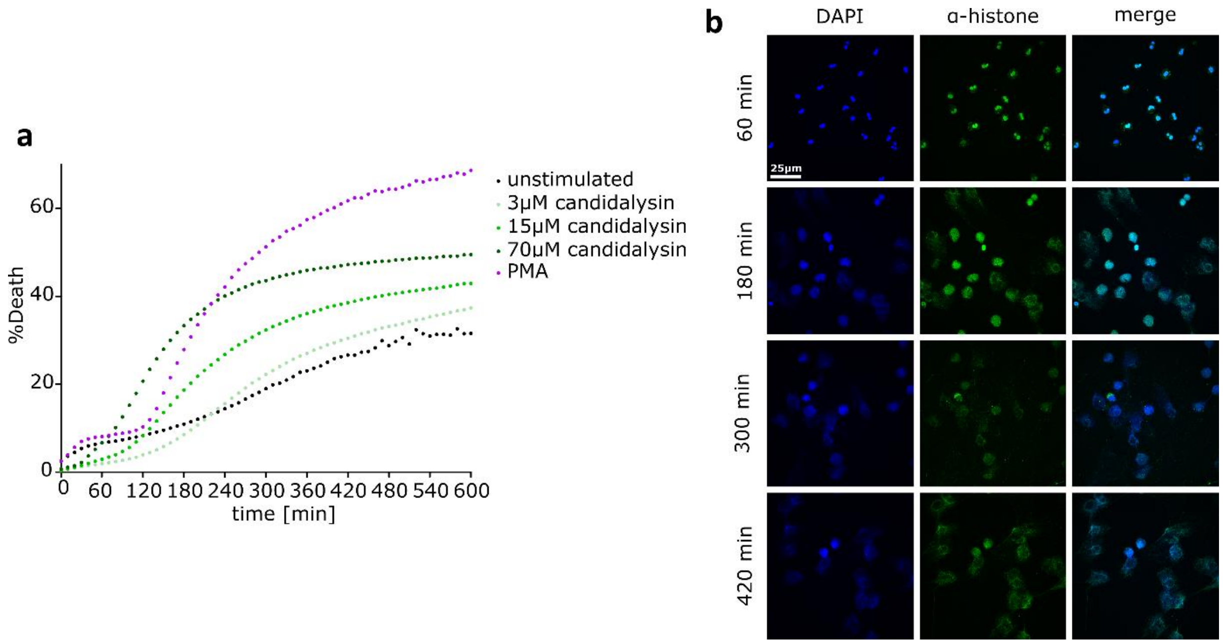


**Fig. S1. Candidalysin and PMA have dose-dependent effects on human neutrophils.** (a) Sytox Green staining demonstrates dose- and time-dependent cytotoxic effects of candidalysin on neutrophils (one representative of 4 donors shown). (b) Immunofluorescent staining of neutrophils treated with 100 mM PMA for 1 h, 3 h, 5 h and 7 h show progressing NET induction. Visualization by DAPI (blue channel) to stain DNA and primary antibody directed against α-histones (green channel). Images taken with Nikon A1R confocal (LSM) controlled by Nikon NIS Elements interface with a Nikon Eclipse Ti-E inverted microscope using 60X magnification.

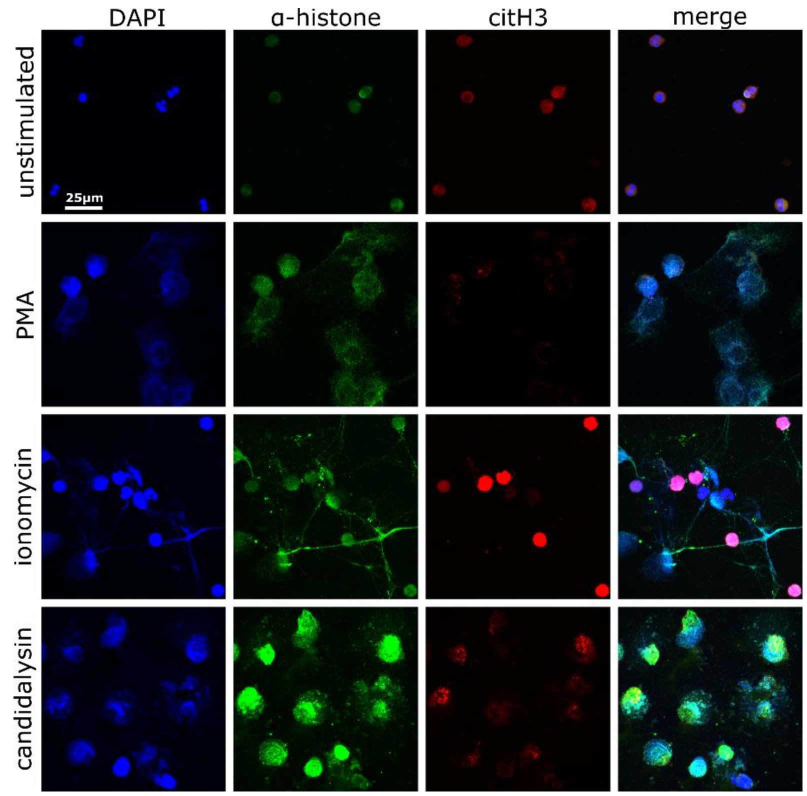


**Fig. S2. Candidalysin induces NET-like structures in human neutrophils.** (a) Immunofluorescent staining of neutrophils treated with 100 nM PMA, 4 µM ionomycin, 1 mM nicotine, 15 µM candidalysin or untreated for 7 h. Visualization by using DAPI to stain DNA (blue channel) and primary antibodies directed against α-histones (green channel) and citrullinated histone H3 (red channel). Images taken with Nikon A1R confocal (LSM) using 60X magnification.

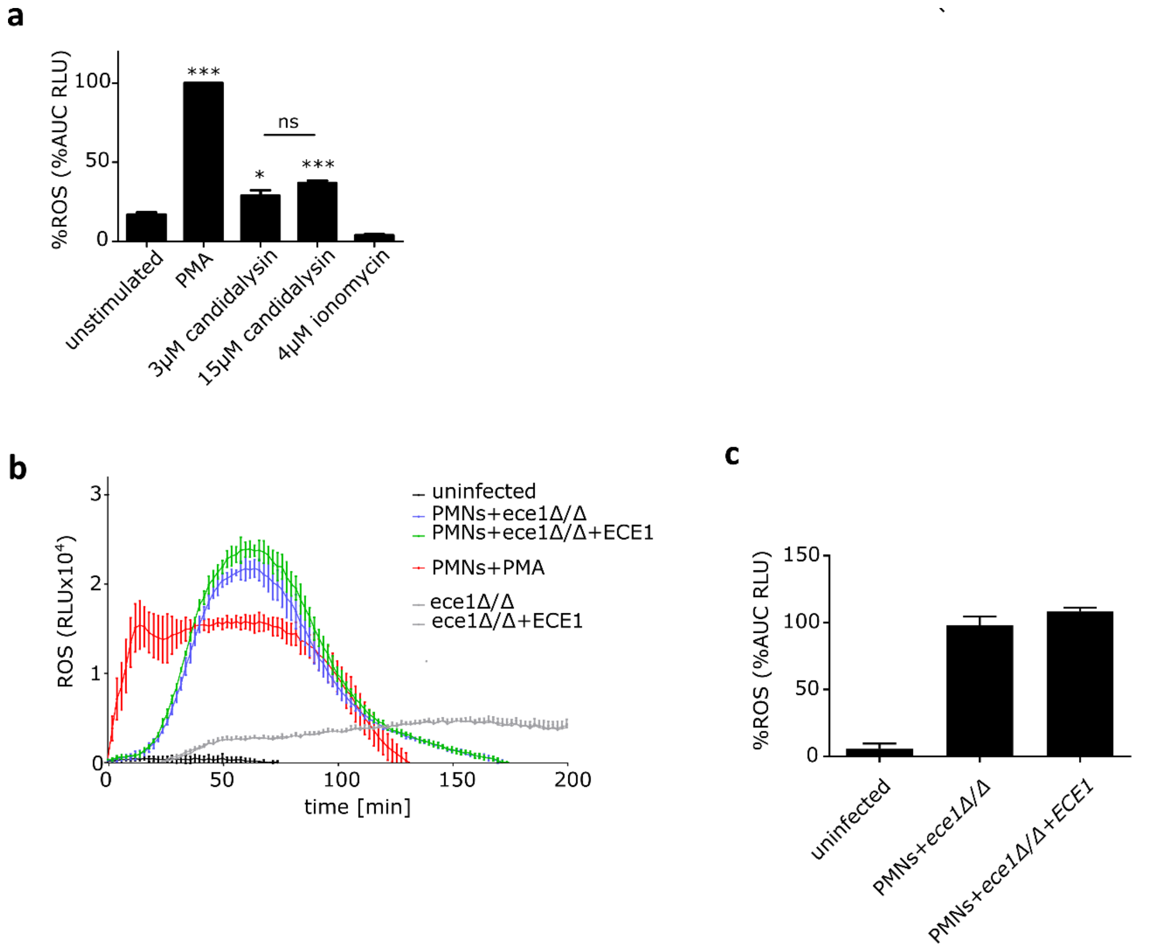


**Fig. S3. Candidalysin triggers measurable ROS response in neutrophils but differences were undetactable in *C. albicans* strains.** ROS production was measured using luminol in (a) PMA, candidalysin and ionomycin stimulated neutrophils over 4 h (n = 3). Infections of neutrophils with *ECE1*$^{-/-}$ *C. albicans* strain showed lower ROS response in comparison to revertant strain displayed as (b) the full time-response of one representative experiment and (c) area under the curve over 3.5 h for 3 combined donors (mean ± SEM). (a,c) represented as to PMA normalized area under the curve. Statistical analysed performed with One-way ANOVA with Bonferroni post-hoc test.

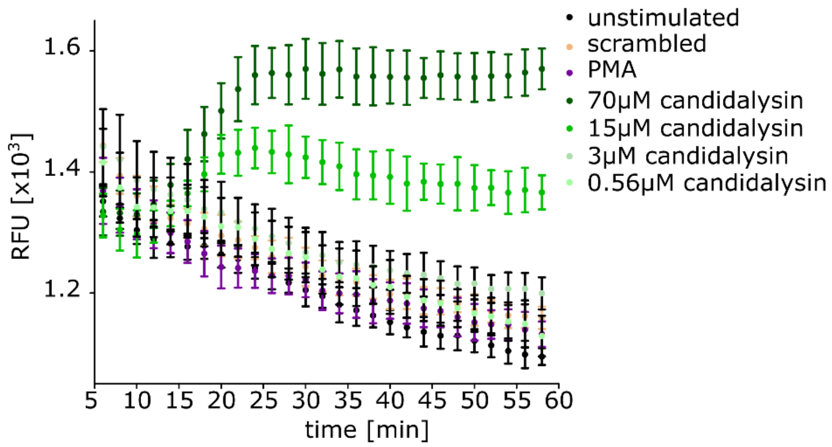


**Fig. S4. Candidalysin causes calcium influx in neutrophils**. Full dose- and time-dependent calcium influx measurement in neutrophils over 1 h induced by candidalysin. Calcium influx was measured with Fluo-8 AM (one representative of 4 donors shown).

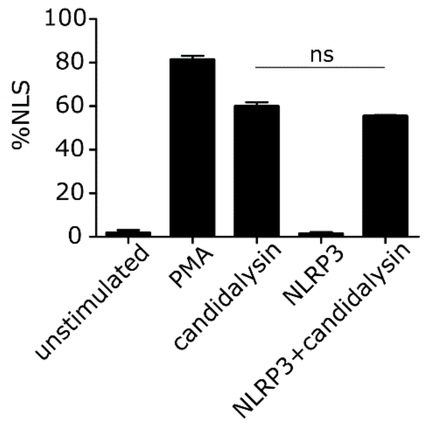


**Fig. S5. NLRP3 inhibition does not affect candidalysin-triggered NLS**. Pharmacological inhibiton of NLRP3 using compound MCC950 (1 µM) did not affect NLS formation in neutrophils after 4.5 h exposure to 15 µM candidalysin. PMA served as positive control (n = 3, mean ± SEM, statistical analysed with one-way ANOVA with Bonferroni post-hoc test).

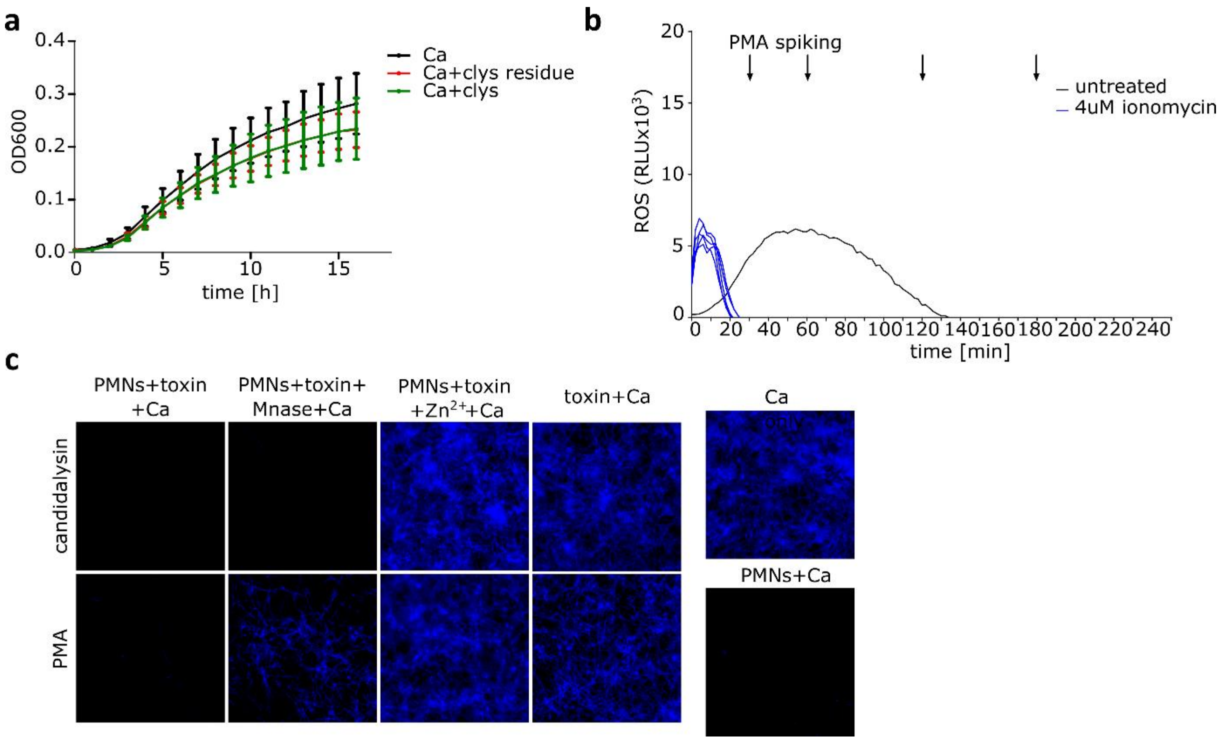


**Fig. S6. Candidalysin-induced NLS but not candidalysin affects fungal growth** . (a) OD measurement confirmed that externally added candidalysin (15 µM) does not inhibit the growth of *C. albicans* (one representative measurement of two biological replicates). (b) Ionomycin treated PMNs showed a minor immiadiate ROS response but were then functionally unable to produce ROS in response to PMA spiking (graph shows one representative of 3 donors). (c) The antimicrobial effect measured through Calcufluor White staining (representative images taken by Cytation 5 Cell Imaging Reader (BioTek) used for quantification in Fig. 8d was caused by NET-like structures alone.
